# Mesenchymal Wnts are required for morphogenetic movements of calvarial osteoblasts during apical expansion

**DOI:** 10.1101/2023.12.05.570300

**Authors:** Nikaya Polsani, Theodora Yung, Evan Thomas, Melissa Phung-Rojas, Isha Gupta, Julie Denker, Xiaotian Feng, Beatriz Ibarra, Sevan Hopyan, Radhika P. Atit

**Affiliations:** Department of Biology and Genome Sciences; Program in Developmental and Stem Cell Biology, Research Institute, The Hospital for Sick Children, Toronto, ON M5G 0A4, Canada; Division of Orthopedics, The Hospital for Sick Children and Departments of Molecular Genetics and Surgery, University of Toronto, Toronto, ON M5G 1X8, Canada; Department of Dermatology and Genome Sciences; Department of Genetics and Genome Sciences

## Abstract

Apical expansion of calvarial osteoblast progenitors from the cranial mesenchyme (CM) above the eye is integral for calvarial growth and enclosure of the brain. The cellular behaviors and signals underlying the morphogenetic process of calvarial expansion are unknown. During apical expansion, we found that mouse calvarial primordia have consistent cellular proliferation, density, and survival with complex tissue scale deformations, raising the possibility that morphogenetic movements underlie expansion. Time lapse light sheet imaging of mouse embryos revealed that calvarial progenitors intercalate in 3D to converge supraorbital arch mesenchyme mediolaterally and extend it apically. In contrast, progenitors located further apically exhibited protrusive and crawling activity. CM cells express non-canonical Wnt/Planar Cell Polarity (PCP) core components and calvarial osteoblasts are bidirectionally polarized. We found non-canonical ligand, *Wnt5a^-/-^* mutants have less dynamic cell rearrangements, protrusive activity, and a flattened head shape. Loss of cranial mesenchyme-restricted *Wntless (*CM-*Wls),* a gene required for secretion of all Wnt ligands, led to diminished apical expansion of OSX^+^ calvarial osteoblasts in the frontal bone primordia in a non-cell autonomous manner without perturbing proliferation or survival. Calvarial osteoblast polarization, progressive cell elongation and enrichment for actin cytoskeleton protein along the baso-apical axis were dependent on CM-Wnts. Thus, CM-Wnts regulate cellular behaviors during calvarial morphogenesis and provide tissue level cues for efficient apical expansion of calvarial osteoblasts. These findings also offer potential insights into the etiologies of calvarial dysplasias.

## INTRODUCTION

The calvaria are a collection of intramembranous bones which form the roof of the skull and protect the brain and other sensory organs. Between each calvarial bone is a mesenchymal suture, which allows for growth and expansion of the skull following birth. Changes in the growth dynamics of calvaria can impact the positions of the sutures and may contribute to congenital calvarial defects such as craniosynostosis and calvarial dysplasias (Flaherty and Richtsmeier, 2018; Miraoui and Marie, 2010; Ornitz and Marie, 2002; Rawlins and Opperman, 2008; Teng et al., 2018; Wilkie and Wall, 1996). These defects are detrimental for brain and sensory organ development (Morriss-Kay and Wilkie, 2005; Wilkie, 1997). Despite the clinical importance of the size and shape of calvaria, we lack a basic understanding of morphogenetic processes by which they are formed. Murine calvarial bones are akin to those of humans, hence the mouse embryo is an ideal system to identify the cellular mechanisms and cues that govern cell movement and apical expansion of mammalian calvarial osteoblast progenitors.

Mammalian calvarial osteoblast progenitors originate from cranial mesenchymal cells derived from cranial neural crest cells (CNCC) and paraxial head mesoderm (PM). The CNCC migrate from the early neural tube alongside the adjacent PM to the supraorbital arch (SOA) beginning by mouse embryonic day (E) 8.5. A subset of this SOA mesenchyme is fated to form the skeletogenic mesenchyme which includes frontal and parietal bone primordia between E10.5-12.5 (Deckelbaum et al., 2012; Tran et al., 2010; Yoshida et al., 2008). At E11.5-12.5, calvarial osteoblast progenitors of frontal and parietal bone primordia begin to condense in foci above the eye and express bone fate transcription factors such as RUNX2. Between E12.5 and 15.5, calvarial osteoblasts commit and express Osterix (OSX/Sp7) while expanding apically (cranially) to cover the brain. While the transcriptional program of mammalian calvarial osteoblast differentiation has been extensively studied (Ang et al., 2022; Fan et al., 2021; Ferguson and Atit, 2018; Ishii et al., 2015), less is understood about the signals and cellular processes that control the morphogenesis and apical expansion of calvarial osteoblast progenitors from the SOA.

Apical expansion and growth of calvaria appears to be a temporally regulated process that can be perturbed by spatio-temporal pharmacological blockade or genetic deletion of various regulatory factors (Ferguson and Atit, 2018; Ferguson et al., 2018; Jiang et al., 2019). Agenesis at the apex of the skull and frontal bone dysplasia is a common phenotype observed following alterations in signaling and transcription factors within the cranial mesenchyme during early calvarial morphogenesis (Ferguson et al., 2018; Goodnough et al., 2016; Jiang et al., 2019; Matsushita et al., 2009; Roybal et al., 2010). For example, the bone transcription program is tightly regulated by several key signaling pathways, including Wnt, BMP, SHH, and FGF signaling which, when dysregulated, often result in defects in calvarial growth (Ang et al., 2022; Fan et al., 2021; Ferguson and Atit, 2018; Ishii et al., 2015). Temporal conditional genetic deletion of transcription factors between E8.5-10.5 such as *Twist1*, *Msx1/2* and epigenetic regulators such as *Ezh2* in cranial mesenchyme result in loss of mineralized bone at the apex of the skull (Dudakovic et al., 2015; Ferguson et al., 2018; Goodnough et al., 2016; Roybal et al., 2010). Since these factors have been associated with apical expansion and growth of calvaria, reduction in expansion may underlie insufficient apical bone formation. However, it is not clear whether calvarial morphogenesis results from various mechanisms regulated by multiple pathways or whether a common downstream cellular mechanism regulates apical expansion.

There are a few putative cellular mechanisms that may contribute to rapid apical expansion of the calvaria (Ishii et al., 2015). One possibility is “end addition” in which cells from ectocranial mesenchyme contribute to the growing osteogenic front under the regulation of *Ephrins* (Ishii et al., 2015; Merrill et al., 2006). However, lineage tracing shows only few cells potentially participate in such a mechanism, suggesting it is likely not the dominant process for apical expansion (Merrill et al., 2006). A second possibility is that spatio-temporally biased proliferation may contribute to apical expansion of calvarial osteoblasts as in other examples of vertebrate morphogenesis(Heisenberg and Bellaïche, 2013; Stooke-Vaughan and Campàs, 2018; Stuckey et al., 2011). In a calvaria *ex-vivo* culture model, inhibition of proliferation lead to a 44% decrease in cell division and decreased growth by only 19%, suggesting that proliferation alone is insufficient to account for apical expansion of maturing calvaria (Lana-Elola et al., 2007). A third potential mechanism is by morphogenetic cell movement such as convergent-extension through cell intercalations as in closure of the neural tube in the frog and fish (Huebner and Wallingford, 2018; Keller et al., 2008; Wallingford and Harland, 2002). Convergent-extension would be consistent with the narrowing of the osteoblast progenitor pool as they progress apically from the SOA. A fourth possibility is directional collective cell migration of calvarial osteoblasts.

There is evidence for long-range displacement of calvarial osteoblasts. When *Engrailed1*-expressing cells were genetically labelled in the SOA at E10.5, they were subsequently found throughout the frontal and parietal bones, among adjacent ectocranial cells, and in the overlying dermis at E16.5 (Deckelbaum et al., 2012; Tran et al., 2010). Consistent with published studies of DiI-labeled mouse calvarial bone primordia at E13.5, the descendants of *En1-*marked cells in the SOA were visible long distances towards the apex (Ting et al., 2009; Yoshida et al., 2008). By both techniques, labeled cells were found much further apically than could readily be accounted for by proliferation alone, strongly supporting the possibility of morphogenetic cell movement during apical expansion of calvarial bones (Tran et al., 2010). However, cellular behaviors and potential tissue-scale cues that might orient efficient expansion of calvarial osteoblast progenitors in the SOA are unknown.

In this study, we found that multiple non-canonical Wnt ligands are expressed in cranial mesenchyme and are required for cellular behaviors engaged in morphogenetic movements and directional migration towards the apex during calvarial morphogenesis. *Wnt5a* contributes to cranial expansion primarily by permitting basal cell rearrangements within the SOA and directional cell movements further apically. This working model provides new insights into the basis of calvarial defects.

## RESULTS

### Two modes of cell movement remodel supraorbital arch mesenchyme

Calvarial osteoblast progenitors that form the murine frontal and parietal bones expand apically from the SOA region between E10.5 to E16.5 (**Fig. 1A**) (Deckelbaum et al., 2012; Tran et al., 2010; Yoshida et al., 2008). Between E13.0 and E14.5, we observed that length-width ratios of frontal bone primordia increased from 5:1 to 10:1 (**Fig. 1B, C**). During this period, a lack of apoptotic cells has been reported within the primordial tissue (**Supp. Fig. 1**), (DiNuoscio and Atit, 2019; Goodnough et al., 2016). To examine the potential role of cell proliferation in this expansion, we tested the proportion of mesenchymal cells in S phase. Uptake of 5-ethynyl-2’-deoxyuridine (EdU) was similar in basal, intermediate and apical regions of the frontal bone primordium at E13.5 and E14.5 (**Supp. Fig. 1 A, B**), an observation that is consistent with data from later stages (Lana-Elola et al., 2007). Therefore, the spatial distribution of proliferating cells does not explain the directional and apically narrowing nature of calvarial expansion. Interestingly, cell shape analysis of E13.5 OSX^+^ calvarial osteoblasts in coronal plane using the membrane marker, Concanvalin, we identified a progressive increase in cellular elongation that corresponded to progressive apical narrowing of the frontal bone primordium (**Fig. 1D**). Spinning disk confocal images of intact embryos show cranial mesenchyme cells in the basal position have protrusive activity that decorate the cell surface (**Fig. 1E)**. In the apical position, cranial mesenchyme cells become elongated and polarized with protrusions present on the apical side of the cells (**Fig. 1F)**. We postulated that dynamic cellular behaviors remodel the SOA mesenchyme as new cells are added.

**Figure 1:**
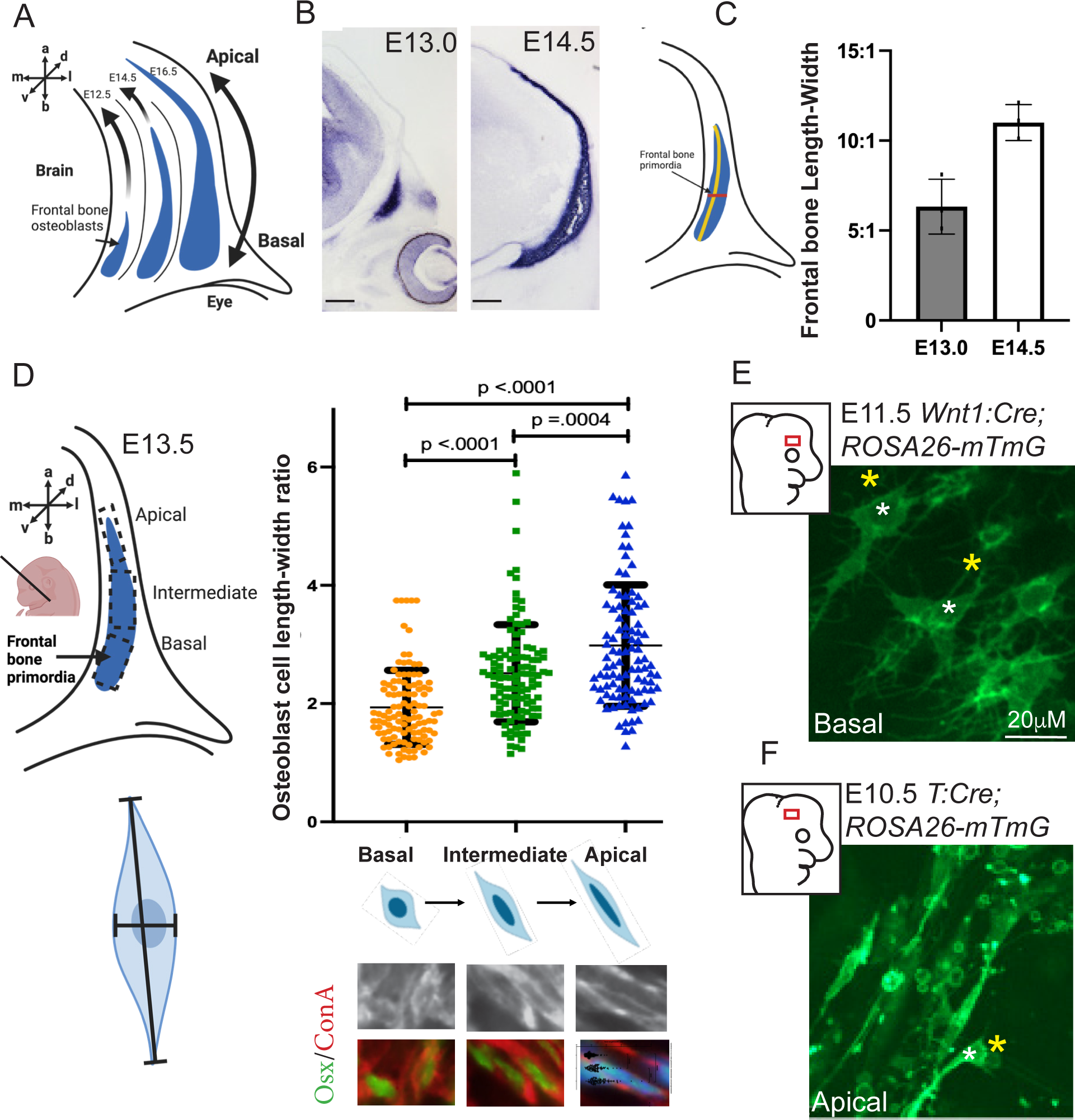
Elongation of frontal bone calvarial osteoblasts occurs at the cellular and tissue level. A) Schematic of the coronal plane view of murine frontal bone primordia expansion from the basal position above the eye towards the apex of the head in embryonic development. B) Alkaline phosphatase (AP) staining of frontal bone primordia (fbp) in the coronal plane at the same magnification. C) Length-width ratio of AP^+^ frontal bone primordia per embryo. D) Schematic of coronal sections indicating position of regions of interest in the fbp for calculating the cellular length-width ratio of E13.5 OSX^+^ calvarial osteoblasts cells (green) co-stained with Concanavalin A (Con A, red) lectin staining for membranes. Significant and progressive increase in cellular elongation from basal to apical position in the fbp. E) Time-lapse confocal imaging of cranial mesenchyme at E11.5 in the basal supraorbital arch (SOA) region showing cells with protrusive activity (yellow asterisk) around the cell body (white asterisk). F) At E10.5 in the apical region, cells are elongated with long protrusions (yellow asterisk) apical to the cell body (white asterisk). Scale bar =100 microns.

To examine dynamic tissue and cell behaviors, we studied an earlier stage of SOA expansion at E10.5 when calvarial osteoblasts precursors are present (Kuroda et al., 2023). Using transgenic embryos that harbor the cell membrane reporter mTmG (Muzumdar et al., 2007) under live conditions (Tao et al., 2019), we applied a strain (deformation) analysis algorithm that we described previously (Tao et al., 2019) and updated (Methods, https://github.com/HopyanLab/Strain_Tool). Briefly, strain was mapped in time-lapse movies by tracking numerous points frame-to-frame and correlating images using a sub-pixel registration algorithm (Guizar-Sicairos et al., 2008) Delauney tessellation was then applied to generate a triangular mesh of the tissue of interest upon which area weighted averages in strain along orthogonal axes were color-coded onto the first frame. In the basal, relatively broad region immediately cranial to the eye, mesenchyme compressed medially and expanded laterally along the mediolateral axis (E_xx_) and extended apically along the apicobasal axis (E_yy_). In contrast, the narrower, more apical mesenchyme predominantly extended along the apicobasal axis. The tissue rotated (reflected by shear - E_xy_) in the basolateral region where it wrapped partially around the orbit and, notably, within the mid-portion of the SOA region mesenchyme between narrowing baso-medial and extending apical regions (**Fig. 2A, Movie 1**). These complex tissue-scale deformations suggest that morphogenetic movements remodel the supraorbital arch.

**Figure 2.**
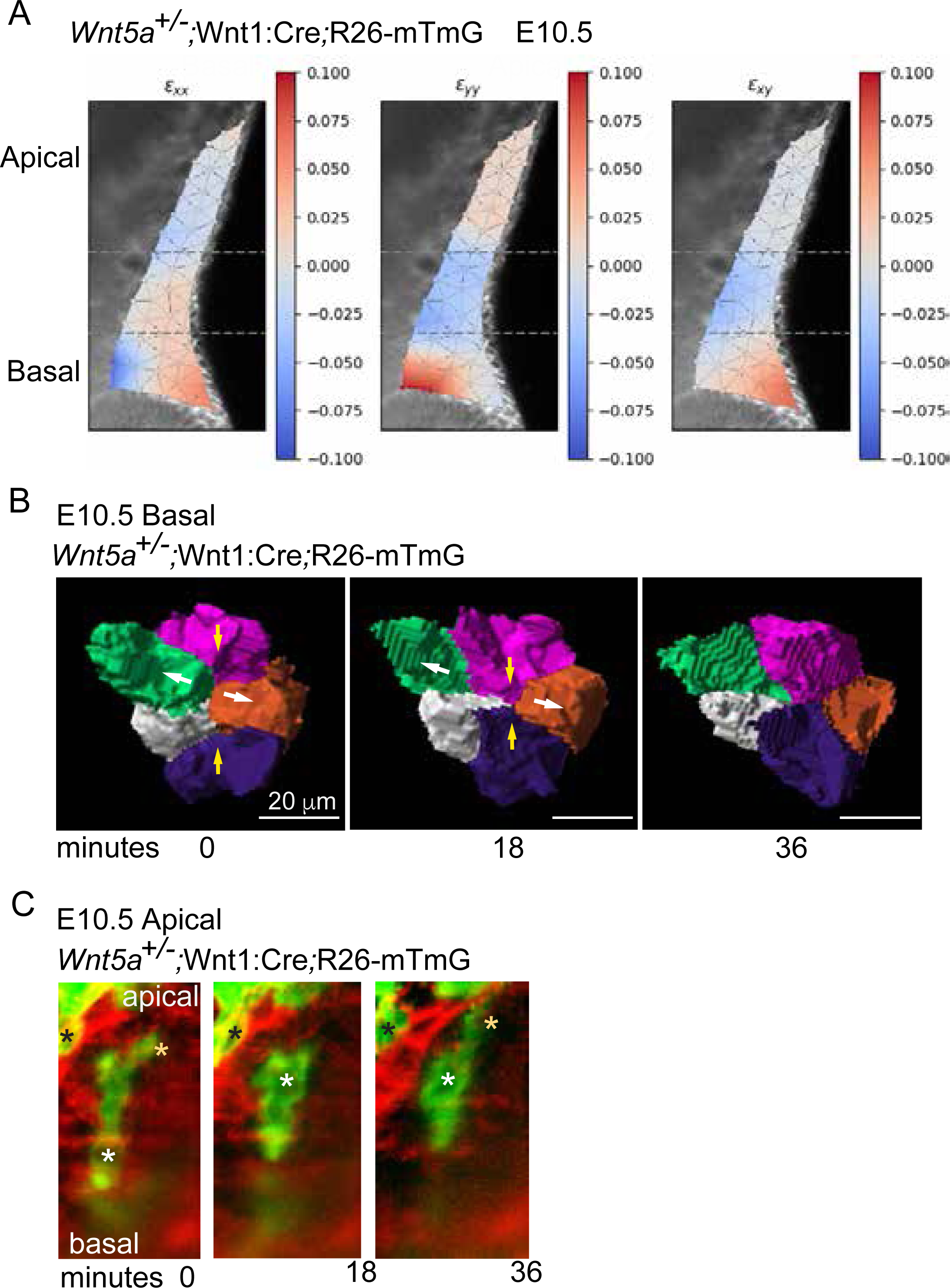
Strain and morphogenetic cell movements in SOA mesenchyme. A) Tissue strain map shown on the first frame of a time-lapse light sheet movie of E10.5 supraorbital arch (SOA) mesenchyme. E*_xx_*, E*_yy_* and E*_xy_* reflect mediolateral, apicobasal and shear, or rotational, deformation, respectively. The colour-coded scale is given as a proportional change (out of 1). B) Time-lapse frames of volume-rendered mesenchymal cells in the basal region of the supraorbital arch of E10.5 *Wnt5a^+/-^*;Wnt1:Cre;R26-mTmG embryos captured via light sheet microscopy. The initial configuration in which green and orange cells are neighbors is changed over 36 min. as they move apart, and magenta and purple cells come together. C) Time-lapse frames of mosaic E10.5 *Wnt5a^+/-^*;Wnt1:Cre;ROSA26-mTmG embryos that allow visualization of cell protrusions. A cell protrusion (yellow asterisk) precedes apical movement of the cell body (white asterisk) with respect to a cell neighbor (black asterisk).

By rendering cell membranes in 3D, we identified mesenchymal cell intercalations within the baso-medial region where convergent extension takes place. Cells exchanged neighbors such that, in the most basic situation, an individual cell entered or exited a surrounding group of about 4-6 other cells (**Fig. 2B, Movie 2**). That process is akin to a 3D T1 exchange that was described for foams (Tao et al., 2019; Weaire et al., 2012) and has been recognized in mandibular arch mesenchyme (Tao et al., 2019). It is not currently possible to track and reliably quantify the relative proportion, complexity, or orientation of these types of 3D intercalations because multiple cells in each identifiable group participate to variable degrees in rearrangements within neighboring groups. Instead, we propose that Golgi positions relative to nuclei offer reasonable proxies for quantification of the spatial distributions of cell orientation (as shown below). We infer that rearrangements among basal cells tend to liquify and permit remodeling of basal mesenchyme.

In contrast, cells within the apical region were elongated and exhibited directional migration. To visualize filipodia that are inserted between neighboring cell membranes, we sparsely converted mTmG-labelled cell membranes from red to green to identify them using a unique *Wnt1:Cre* strain that has become mosaically expressed in our colony. Cell protrusions oriented toward the apex or base of the SOA preceded movement of the cell body in the same direction (**Fig. 2C, Mov. 3, 4**). Based on their elongated shapes (**Fig. 1D**), these bipolar cell movements should tend to elongate mesenchyme along the apicobasal axis. A transition of cellular behaviors from basal rearrangements to apical shear-type movements contributes to remodeling of the SOA.

### Mesenchymal non-canonical Wnts are required for apical expansion of the frontal bone primordium

Non-canonical Wnt pathway and Wnt5a/11 ligands are key regulators of cell polarity, convergent-extension, and directional cell movement (Butler and Wallingford, 2017; Gray et al., 2011; Zallen, 2007). After modifications by *Wntless*, Wnt ligands are secreted from the cell and bind to cell surface Frizzled receptors and co-receptors, ROR2 and VANGL2 to initiate a signaling cascade (Andre et al., 2015; Davey and Moens, 2017). We observed robust expression of key signaling components *Wnt5a*, *Wnt11*(Goodnough et al., 2014), the co-receptor ROR2, and phosphorylated c-Jun, a downstream effector of non-canonical Wnt signaling, throughout the cranial mesenchyme between E11.5-13.5 (**Supp. Fig. 2A-I**). Interestingly at E11.5, VANGL2, a regulator of cytoskeleton-mediated changes in cell polarity, was readily identifiable by immunostaining in basal, but not apical, mesenchyme suggesting pathway function, like cell behaviors, is different in those two regions (**Supp. Fig. 2 F, G**). Overall, these data suggest that non-canonical Wnt/PCP signaling is active within cranial mesenchyme during the remodeling of SOA.

To test for Wnt/PCP pathway function, we conditionally deleted mesenchymal *Wntless* to prevent secretion of all Wnt ligands and circumvent functional redundancy. We used tamoxifen-inducible *Pdgfrα:Cre-ER* to delete *Wntless* in cranial mesenchyme (CM-*Wls^fl/fl^*) (alternative: *Pdgfrα:Cre-ER; Wls^fl/fl^*) **(Fig. 3A)**. Since Cre-recombination occurs within 24 hours after oral gavage that was initiated at E8.5, we obtained efficient deletion of *Wls* mRNA by E12.5 (DiNuoscio and Atit, 2019; Ibarra et al., 2021). With this method, the cranial mesenchyme loses its ability to secrete Wnt ligands; however, the ectodermal Wnts are sufficient to maintain canonical Wnt pathway activation through E12.5 (DiNuoscio and Atit, 2019; Ibarra et al., 2021) **(Fig. 3A)**. Fate mapping of *PdgfraCre*-ER expressing cells between E9.0-10.5 showed β-galactosidase^+^ cells were present broadly in the cranial mesenchyme including the in the frontal bone primordia and adjacent dermis in controls and in CM-*Wls^fl/fl^* mutants at E14.5 **(Fig. 3B, C)**.

**Figure 3:**
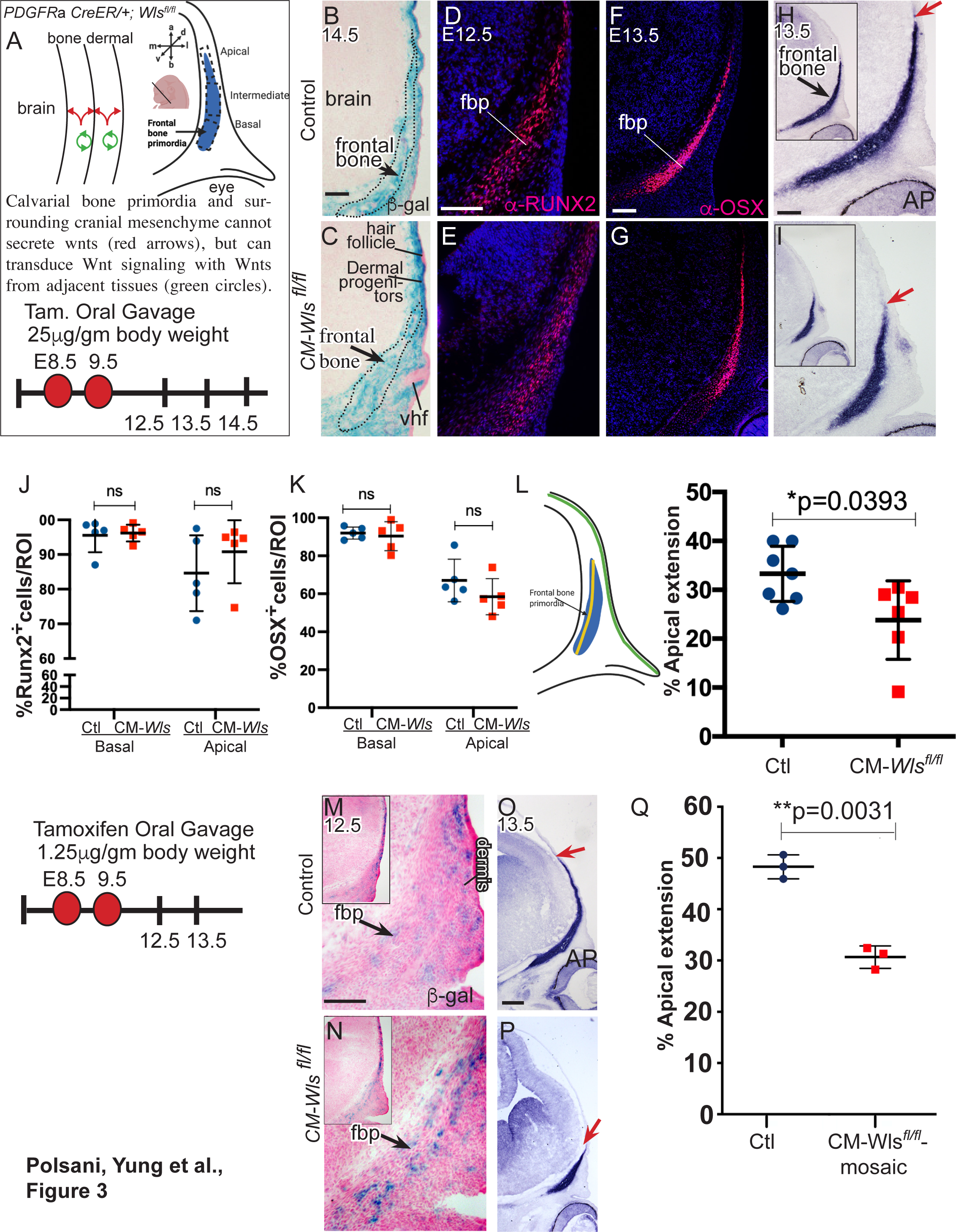
Mesenchyme Wnts are required for efficient expansion of frontal bone primordia in a non-cell autonomous manner. A) Schematic of the conditional and inducible deletion of *Wntless* in SOA cranial mesenchyme (CM-*Wls)* mouse embryo. B,C) Lineage labeled R26R-β-galactosidase (blue) expression in the SOA region cranial mesenchyme in controls and CM-*Wls* mutants. D-I) Protein expression of RUNX2, OSX by indirect immunofluorescence and Alkaline Phosphatase (AP) expression in the frontal bone primordia (fbp). J-K) Percent RUNX2^+^ cells at E12.5 and OSX^+^ at E13.5 in the basal and apical region of the fbp in regions of interest (ROI) as indicated in panel (A). L) Schematic for calculating the percent AP^+^ frontal bone apical extension (yellow line) and normalized to cranial length (green line). Apical extension of fbp is significantly lower in CM-*Wls* mutants. M-N) Low dose tamoxifen regimen and recombination of R26R-β-gal reporter showing mosaic labeling in the SOA mesenchyme in control and CM-*Wls* mutants. O-Q) Mosaic deletion of CM-*Wls* leads to significant decrease in normalized apical expansion of fbp. H,I, O, P) Apical extent of the fbp is marked by red arrows. Scale bar =100 microns.

At E14.5 and 16.5, CM-*Wls^fl/fl^* mutants exhibited diminished expansion of the forelimb and craniofacial structures, and a curly tail, phenocopying mutants of other core-PCP components (Gao et al., 2011) (**Supp. Fig. 4A-D**). CM-*Wls^fl/fl^* mutants expressed the early calvarial osteoblast markers, RUNX2, OSX, and alkaline phosphatase (AP) (**Fig. 3B-I**), indicating that mesenchymal Wnts are not required for cell fate specification and early commitment to the osteoblast lineage. The proportion of RUNX2 and OSX expressing cells per region of interest was comparable in the basal and apical portion of the frontal bone primordia **(Fig.3J, K)**. In contrast, ectodermal Wnt-dependent activation of mesenchymal canonical Wnt/β-catenin signaling is required for osteoblast differentiation (Goodnough et al., 2012). Thus, we infer that the CM-*Wls* mutant phenotype here is primarily due to a lack of noncanonical Wnt signaling.

The normalized apical extension and area of AP-positive frontal bone primordia was significantly diminished among CM-*Wls^fl/fl^* mutants relative to controls at E13.5 **(Fig. 3L, Supp Fig. 3C-E)**. The decrease in apical expansion of frontal bone primordia in CM-*Wls^fl/fl^* mutant embryos was not attributable to cell proliferation index, cell density, or cell survival which were comparable to the controls (n= 4-7, 2-3 liters) (**Supp Fig. 3F-J**).

The versatility of the conditional, inducible *Pdgfrα:Cre-ER* allowed us to diminish the extent of Cre-recombination within cranial mesenchyme. By significantly decreasing the dose of tamoxifen (by approximately 25-fold) at E8.5, we limited the number of cells in which *Cre* was activated, effectively generating a mosaic population of mutants. We used R26R β-gal to monitor Cre-ER activity within the CM starting at E12.5 **(Fig 3M, N)**. Mosaic loss of *Wls* in a small number of mesenchymal cells was sufficient to attenuate apical expansion of frontal bone primordia as measured by AP-staining **(Fig 3O-Q)**. Next, we used the *PDGFRα:Cre-ER*;R26 tdTomato reporter to monitor the distribution of Cre-ER^+^ descendants. Similar to control embryos, tdTomato^+^ cells were distributed throughout the OSX+ frontal bone primordium, in the adjacent dermal mesenchyme, and cranial mesenchyme that is further apical to the calvarial osteoblast domain at E13.5 **(Supp to Fig. 4 A-E)**. Together, these data indicate that mesenchyme *Wls* is not required for osteoblast commitment to the bone lineage but for apical expansion of the frontal bone primordium in a non-cell autonomous manner.

### Wnt5a and CM-Wnts facilitate morphogenetic cell movements

The preceding data suggest that *Wls* regulates paracrine cues that affect cellular parameters to remodel SOA mesenchyme. To study calvarial osteoblasts cellular shapes in mutants, we examined the nuclear length-width ratio of calvarial osteoblasts because it is feasible and consistently tracks cell shape (Chen et al., 2015; Green, 2022) **(Fig. 1D)**. Runx2^+^ nuclei shape elongation was reduced, and circularity was increased in the basal and apical regions of mutant embryos **(Fig. 4A, B)**. The enrichment of filamentous actin in the cranial mesenchyme apically, as assessed by phalloidin staining intensity, was also diminished **(Fig. 4C-E)**, indicating CM-Wnts contribute to cellular elongation and cytoskeletal organization.

**Figure 4:**
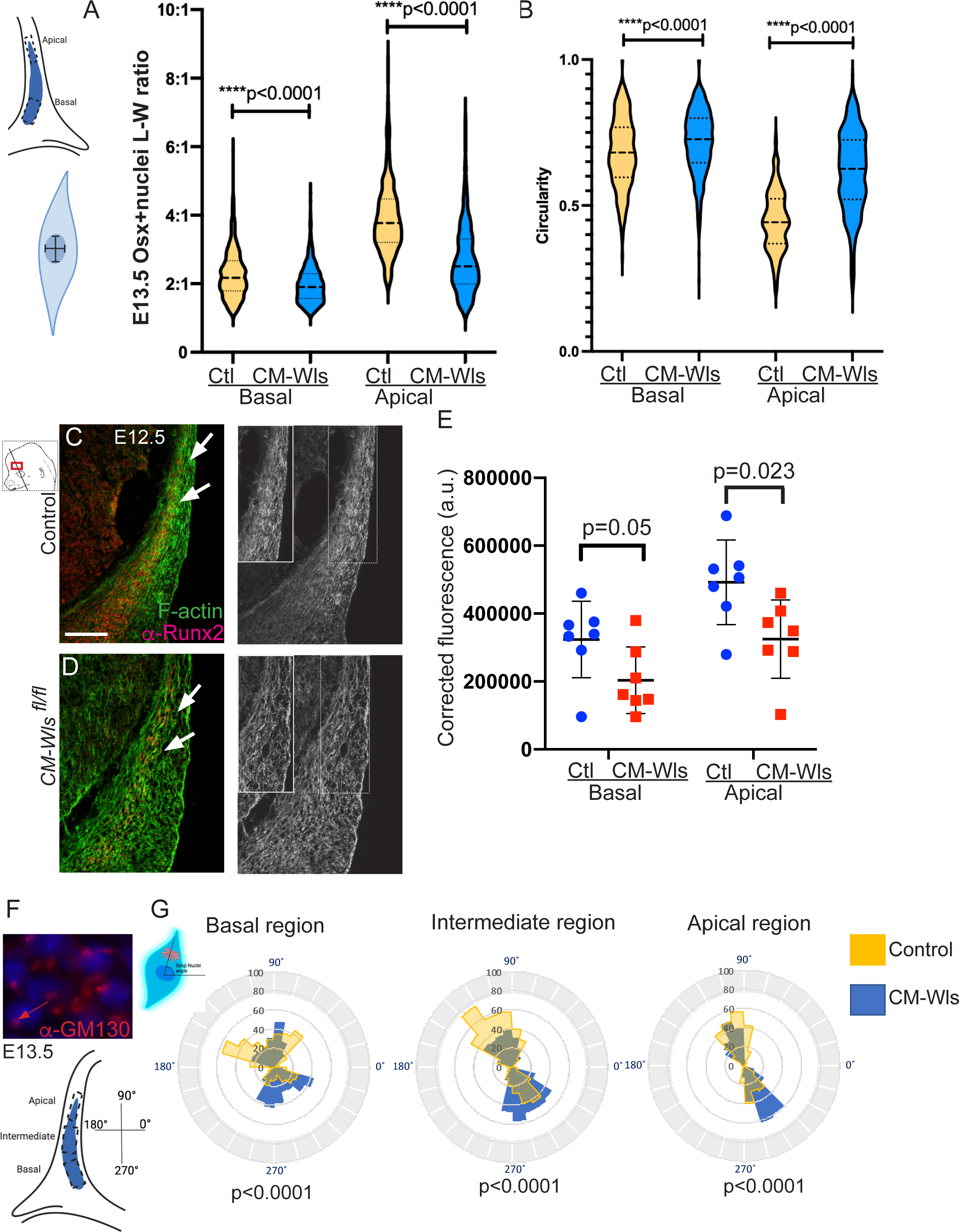
Calvarial osteoblasts cellular behavior associated with migration is dependent on mesenchyme Wnts. A,B) Nuclei shape analysis showing a significant decrease in length-width ratio and consistently increase in circularity in OSX^+^ calvarial osteoblasts at E13.5. C-D) Filamentous actin (F-actin) stained with phalloidin shows enrichment in apical region of the RUNX2^+^ frontal bone primordia (fbp) and the apical cranial mesenchyme at E12.5 in the controls that is diminished in *CM-Wls* mutants (white arrows). E) Corrective fluorescence of F-actin expression in the basal and apical region of fbp showing a significant decrease in enrichment of F-actin in the CM-*Wls* mutants. F) Calvarial osteoblasts polarization was visualized by measuring the GM130^+^ Golgi complex (red)-DAPI+ nuclei angle as shown in the schematic in the ROIs of the fbp at E13.5. G) In controls, the Golgi complex-nuclei angle indicate calvarial osteoblasts are bidirectional and progressively more polarized along the axis of growth. The distribution of Golgi complex-nuclei angle is significantly altered in each region of the fbp in *CM-Wls* mutants. Scale bar =100 microns.

Since the positions of Golgi with respect to nuclei correspond to axes of cell polarity (Ede and Wilby, 1981; Yadav et al., 2009), we measured GM130^+^ Golgi-nuclei angles among calvarial osteoblasts within the OSX^+^ domain of E13.5 frontal bone primordia. With respect to the midline of the brain as a reference axis, Golgi-nuclei angles were distributed more broadly in the basal region than the apical region **(Fig. 4F,G)**. In the intermediate and apical regions, wild-type and mutant cells were increasingly bi-polar along the apical-basal axis. Comparatively, more mutant cells exhibited a basal-ward bias in all regions and the distribution of Golgi-nuclei angles were significantly different in *CM-Wls* mutants in each region (**Fig. 4F,G)**. These data indicate that cell polarity is progressively biased along the apical-basal axis of growth and the repositioning of the Golgi complex to the front of the nuclei is dependent upon CM-*Wls*.

A likely paracrine effector in the SOA mesenchyme is WNT5A (**Supp Fig.3)**. Using *Wnt5a^-/-^* embryos labelled with the transgenic membrane reporter mTmG, convergent-extension-type remodeling of the basal tissue and extension of apical tissue were diminished by time-lapse light sheet microscopy and strain analysis **(Fig. 5A, Mov. 5)**. At the basal aspect of the mutant SOA, mean strain was similar to that of wild-type embryos **(Fig. 5B)**. However, the variance from the mean (root mean square) of strain among individual foci was diminished in mutants (E_xx_ root mean square *Wnt5^+/-^* = 0.18, *Wnt5a^-/-^* = 0.12, t test p = 0.02, n = 3 embryos; E_yy_ root mean square *Wnt5^+/-^* = 0.18, *Wnt5a^-/-^* = 0.13, t test p = 0.03, n = 3 embryos), indicating that cell rearrangements were less frequent or complete. Consistent with this possibility and the altered Golgi positions among mutant cells (**Fig. 4G**), time-lapse imaging revealed a paucity of cell rearrangements in 3D. To better illustrate this qualitative observation, we include 3D rendered movies from 3 separate embryos within the basal SOA. In those movies, cell shapes fluctuate but outright neighbor exchange is not observed **(Fig. 5C, Supp Fig. 5A, B, Mov 6-8)**. In apical mesenchyme, mediolateral narrowing (E_xx_) and apicobasal extension (E_xy_) were diminished among *Wnt5a^-/-^* embryos. Consistent with this finding and with altered cell orientation based on Golgi positions (**Fig. 4G**), protrusive activity among apically located mutant cells was qualitatively less abundant and multidirectional, rather than bipolar **(Fig. 5D, Mov. 9)**. We conclude that *Wnt5a* is required to remodel the SOA mesenchyme by facilitating morphogenetic cell movements that restrain mediolateral expansion of the base while promoting elongation of the apical aspect of the SOA mesenchyme.

**Figure 5.**
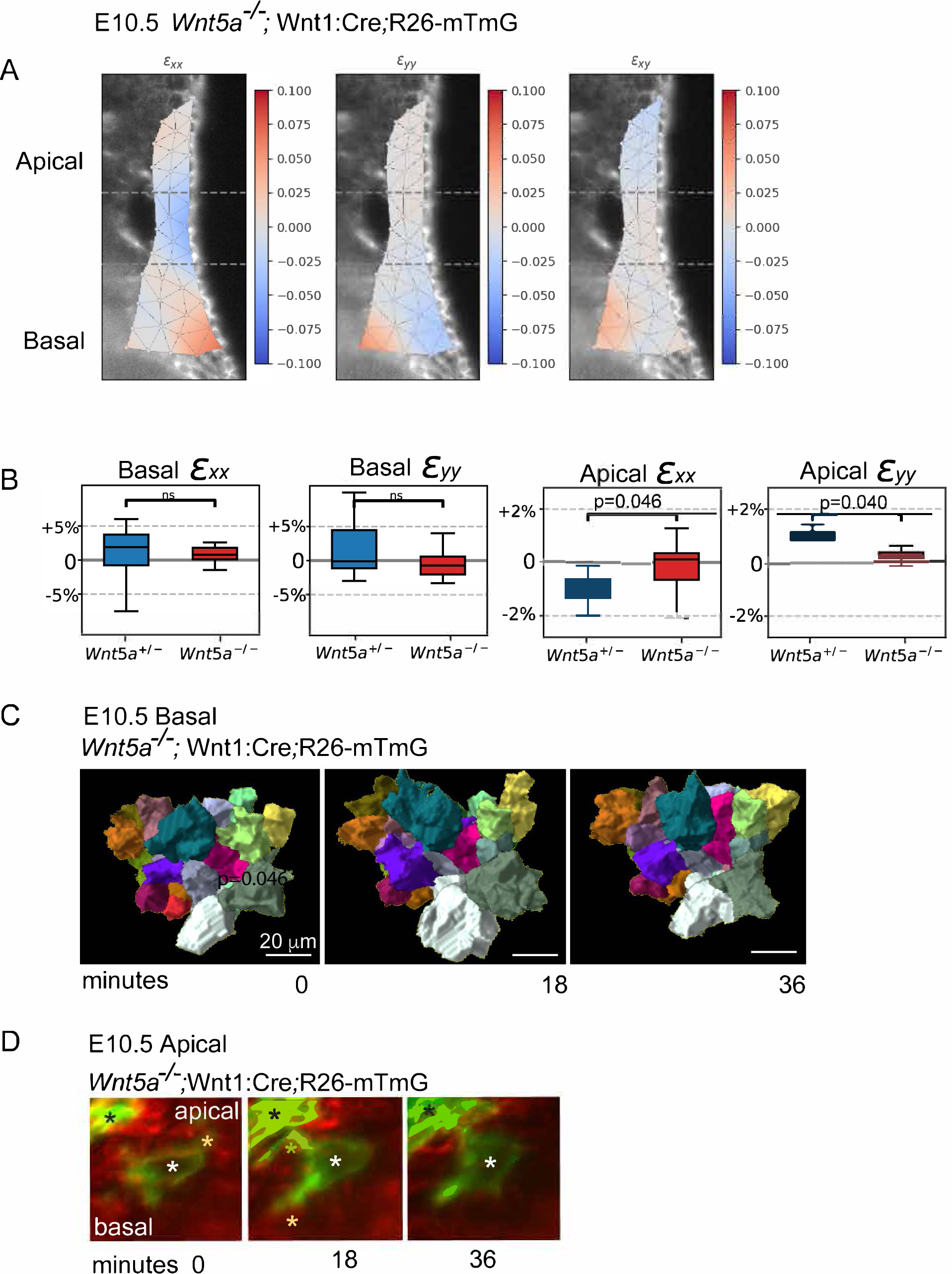
Impaired cell rearrangements in *Wnt5a^-/-^*mutants. A) Tissue strain map shown on the first frame of a time-lapse light sheet movie of E10.5 *Wnt5a^-/-^* supraorbital arch mesenchyme. E*_xx_*, E*_yy_* and E*_xy_* reflect mediolateral, apicobasal and shear, or rotational, deformation, respectively. The color-coded scale is given as a proportional change (out of 1). B) Quantification of tissue strain in control and *Wnt5a^-/-^* mutant. In the basal region, the mean change is similar between genotypes but deviation from the mean is greater among WT embryos (E_xx_ root mean square *Wnt5a^+/-^*=0.18, *Wnt5a^-/-^*=0.12, t test p=0.02, n=3 embryos; E_yy_ root mean square *Wnt5a^+/-^*=0.18, *Wnt5a^-/^* =0.13, t test p=0.03, n=3 embryos), suggesting dynamic cell behaviors may be diminished in the mutants. In apical mesenchyme, mediolateral narrowing (E_xx_) and apicobasal extension (E_yy_) were diminished among *Wnt5a^-/-^* embryos. B) Time-lapse frames of volume-rendered mesenchymal cells in the basal region of the supraorbital arch of E10.5 *Wnt5a^-/-^;* Wnt1:Cre;R26:mTmG embryos captured via light sheet microscopy. Despite cell shape fluctuations, neighbor positions remain stable. C) Time-lapse frames of mosaic E10.5 *Wnt5a^-/-^;*Wnt1:Cre;R26:mTmG embryos that allow visualization of cell protrusions. D) Protrusive activity (yellow asterisks) is present and multidirectional. The protrusive cell body (white asterisk) does not progress relative to a cell neighbor (black asterisk).

Collectively, our work reveals calvarial osteoblasts exhibit two different modes of cell movements and complex cellular behaviors that are regionalized along the baso-apical axis during calvarial morphogenesis *in vivo*. Mechanistically, we found these behaviors such as cell intercalation, cellular elongation, protrusive activity, and filamentous actin enrichment are achieved via non canonical cranial mesenchyme Wnts and in particular Wnt5a (**Figure Supplementary 6**). Thus, our data improves upon our understanding of the physiological and pathological processes of directional expansion in mammalian morphogenesis and its involvement calvarial defects.

## DISCUSSION

Despite the even distribution of mitotic cells within the calvarial primordia, remodeling during growth is achieved by cell movements. Other examples of this type of growth control in mouse embryonic mesenchyme include the limb bud (Boehm et al., 2010; Gros et al., 2010; Wyngaarden et al., 2010)and mandibular arch (Tao et al., 2019), and epithelial examples include the anterior visceral endoderm (Trichas et al., 2012) and kidney tubules (Lienkamp et al., 2012). The transition of basal cell rearrangements to directional apical cell movements underscores how, as morphogenetic mechanisms alter tissue shape, different intercellular strategies are employed.

Requirement for the noncanonical Wnt pathway in a paracrine fashion for two distinct modes of cell movement *in vivo* in the adjacent basal and apical region of SOA mesenchyme is interesting. It confirms the recognized role of the non-canonical Wnt pathway in facilitating cytoskeletal organization and contractions (Davey and Moens, 2017; Shindo, 2018; Shindo et al., 2019). However, our data imply the pathway does not necessarily control the type or orientation of contraction. Cell shape oscillations that drive junctional rearrangements (Gorfinkiel, 2016; Levayer and Lecuit, 2013; Tao et al., 2019) and protrusive activity that drives directional migration (Carmona-Fontaine et al., 2008) represent overlapping yet distinct cytoskeletal activities (Huebner and Wallingford, 2018). Although the SOA and frontal bone primordia does not elongate normally in CM-*Wls* mutants, our evidence suggests that result is not due to the loss of a Wnt orientation cue. Consistent with the broad distribution of the Wnt5a and Wnt11 ligands in the cranial mesenchyme, neither basal cells nor apical cells were unidirectionally oriented in control embryos. Furthermore, loss of *Wnt5a* diminished but did not abolish both the basal junctional rearrangements and the bipolar nature of the more apically located cells and protrusive activity. Those observations raise the possibility that the noncanonical Wnt pathway is permissive for morphogenetic cell movements and that other factors, such as tissue-scale geometric constraint or forces (Lau et al., 2015; Yamada and Sixt, 2019), physical (Aigouy et al., 2010; Shellard and Mayor, 2021; Zhu et al., 2020) and/or other extracellular cues (Koca et al., 2022; Yamada et al., 2022), potentially orient tissue growth.

We show that morphogenetic cell movements *in vivo* and associated cellular behaviors are new players to calvarial bone morphogenesis and apical expansion. The role of cell movement and multiple modes of cell movement in craniosynostosis and congenital calvarial dysplasias remains to be investigated and can advance our morphogenetic understanding of congenital defects.

## Movie Legends

**Movie 1.** Light sheet time lapse movie of the supraorbital arch of an intact mouse embryo that harbors a transgenic fluorescent cell membrane label (mTmG) at E10.5. The orbit is apparent at the bottom of the field. This coronal view of a mediolateral (left-to-right) and apicobasal (top-bottom) plane shows evidence of tissue remodeling with basal narrowing and apical extension.

**Movie 2.** A group of cell neighbors rendered in 3D and pseudo-colored that were within the basal region of the supraorbital arch of an intact mTmG mouse embryo at E10.5. Highly dynamic cell shape changes occur as cell neighbor exchanges take place. The green and white cells lose their junctions with the orange cell as the pink and purple cells make contact over 1.5 h.

**Movie 3.** Time lapse light sheet movie of a Wnt1:Cre;mTmG embryo at E10.5. The field of view is within the apical region of supraorbital mesenchyme near the adjacent lateral ectoderm on the left. Cell membranes are sparsely labelled in green due to mosaicism of the Cre allowing for visualization of filopodia. Filopodial extension precedes movement of the cell body (dark) toward the apex.

**Movie 4.** Time lapse light sheet movie of a Wnt1:Cre;mTmG embryo at E10.5. The embryo is different to the one shown in Movie 3. The field of view is within the apical region of supraorbital mesenchyme near the adjacent lateral ectoderm on the left. Cell membranes are sparsely labelled in green due to mosaicism of the Cre allowing for visualization of filopodia. Filopodial extension precedes apical movement of the cell body (dark) just deep to the ectoderm.

**Movie 5.** Time lapse light sheet movie of the supraorbital arch of a *Wnt5a^-/-^*; mTmG embryo at E10.5. Visual evidence of basal mediolateral narrowing and apical extension is diminished relative to the wild-type condition (Movie 1).

**Movies 6-8.** Time lapse light sheet movies of the basal supraorbital regions of three separate *Wnt5a^-/-^*; mTmG embryos at E10.5. These 3D renderings demonstrate cell shape fluctuations but cell neighbor exchange that is easily seen in wild-type embryos (Movie 2) is not readily identifiable in the mutants.

**Movie 9.** Time lapse light sheet movie of a *Wnt5a^-/-^*;Wnt1:Cre;mTmG embryo at E10.5. The field of view is within the apical region of supraorbital mesenchyme near the adjacent lateral ectoderm on the left. Cell membranes are sparsely labelled in green due to mosaicism of the Cre allowing for visualization of filopodia. Filopodial activity is present but not oriented apically or basally and the cell body (dark) moves little in response to the protrusions.

**Supplemental Figure 1:**
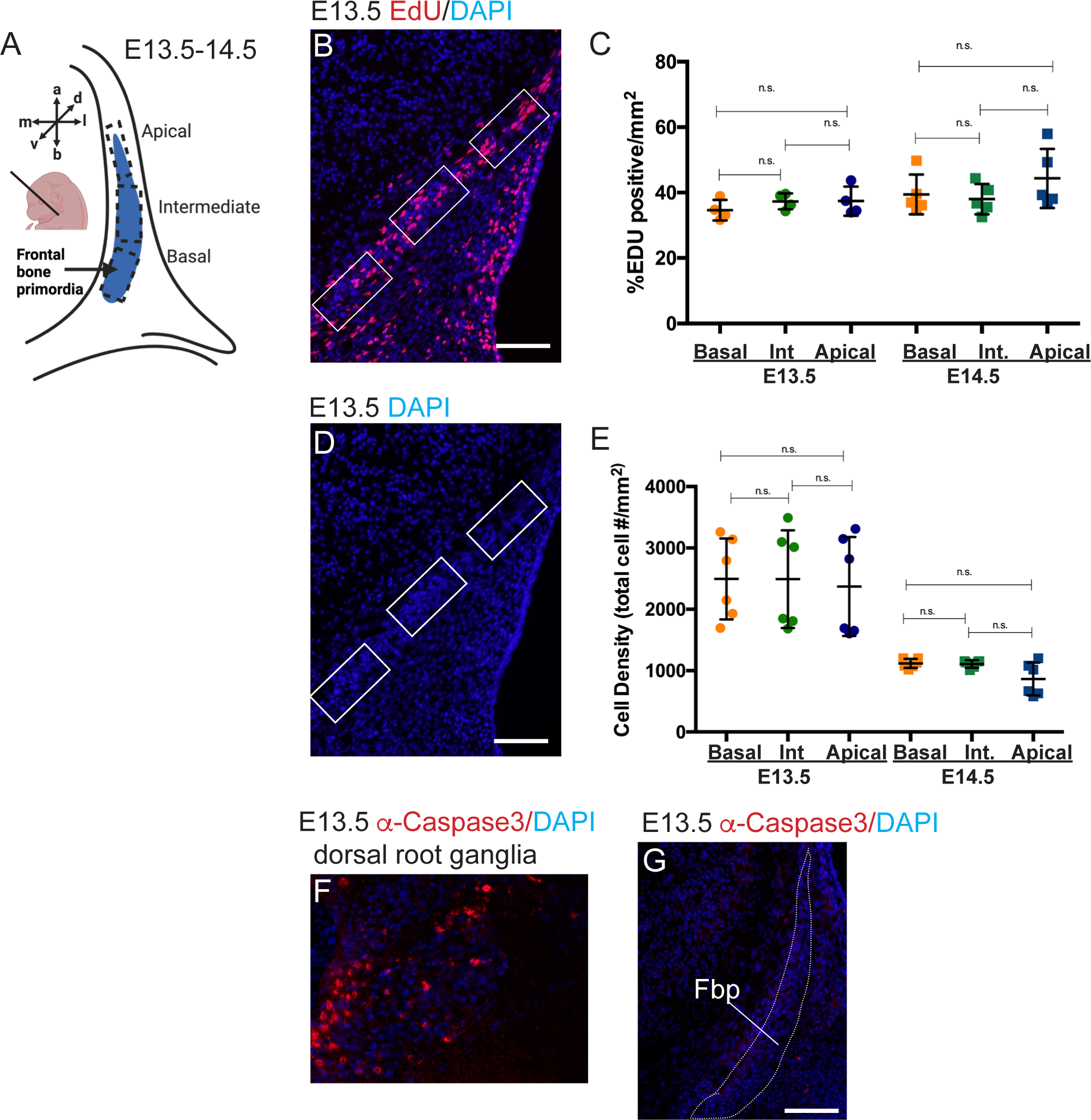
Proliferation and cell survival in control frontal bone primordia during apical expansion. A,B) Schematic of the coronal section showing regions of interest (ROI) in the basal, intermediate and apical portion of the frontal bone primordia (fbp) stained for EdU incorporation. C) Proliferation index was calculated from percent EdU^+^ nuclei/total DAPI^+^ nuclei per ROI in the fbp. D, E) Cell density was calculated by number of DAPI^+^ nuclei per ROI in the fbp. F-G) Cell survival was visualized by expression of activated caspase 3 protein. Positive control region in the dorsal root ganglia. Scale bar =100 microns

**Supplemental Figure 2:**
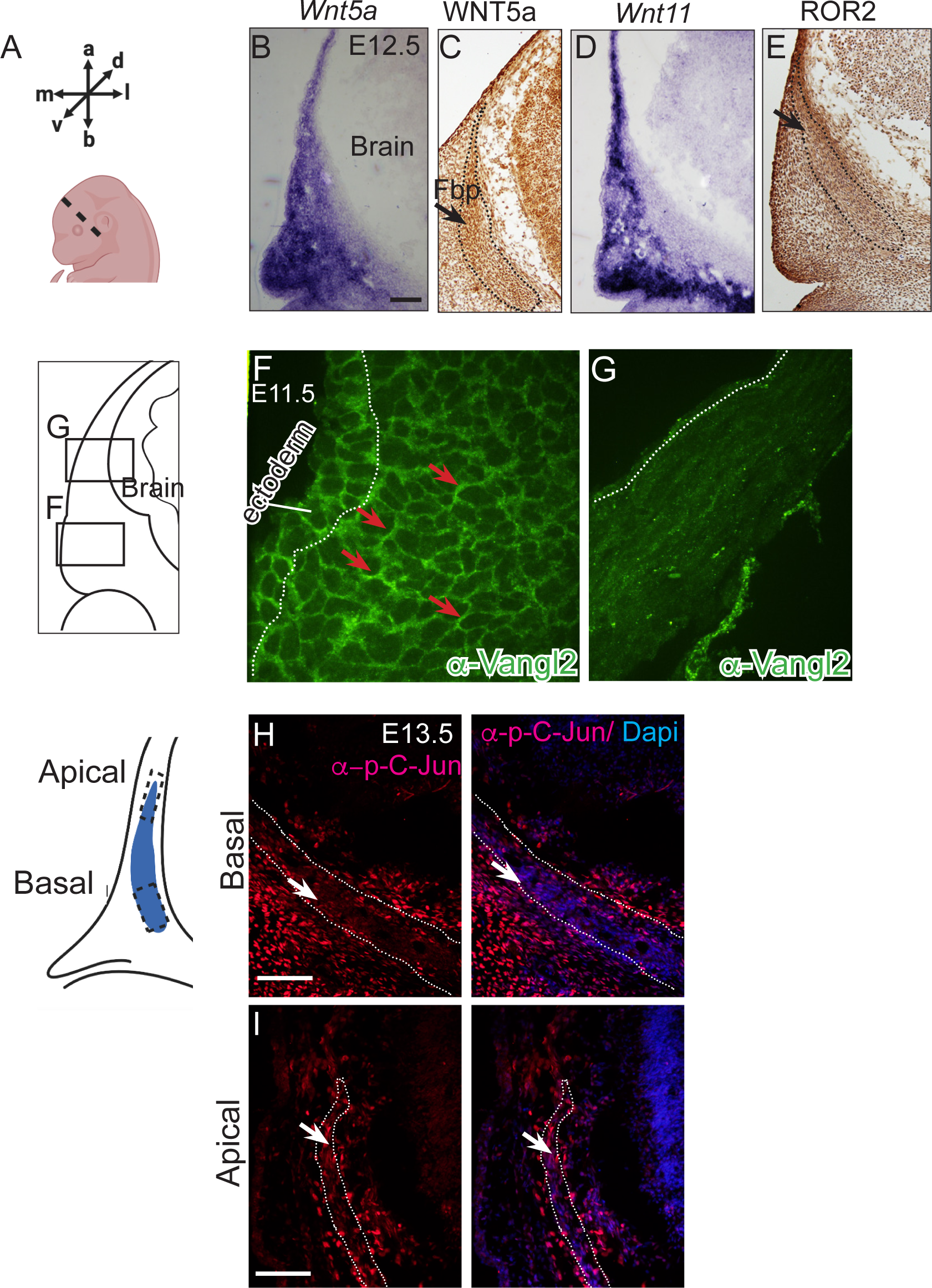
Expression of core components of non-canonical Wnt/PCP pathway in the SOA mesenchyme. A) Schematic of the frontal bone primordia in the coronal plane at E12.5-13.5 showing regions of interest in the basal and apical position. B-E) Expression of non-canonical Wnt pathway ligands Wnt5a mRNA and protein and Wnt11 and ROR2 receptor is expressed broadly in the SOA mesenchyme containing frontal bone primordia (Fpb, black dotted lines, black arrow). F-G) Expression of VANGL2 protein by indirect immunofluorescence shows asymmetric expression in cells in basal region (red arrows) of the SOA mesenchyme and ectoderm and absent in the apical region of the cranial mesenchyme at E11.5. H-I) Expression of phosphorylated c-Jun/JNK by indirect immunofluorescence is visible in the cranial mesenchyme in the periphery of the fbp in the basal region and in the apical region at E13.5 (white arrow), indicating regional activation of non-canonical Wnt/PCP signaling. Scale bar =100 microns

**Supplemental Figure 3:**
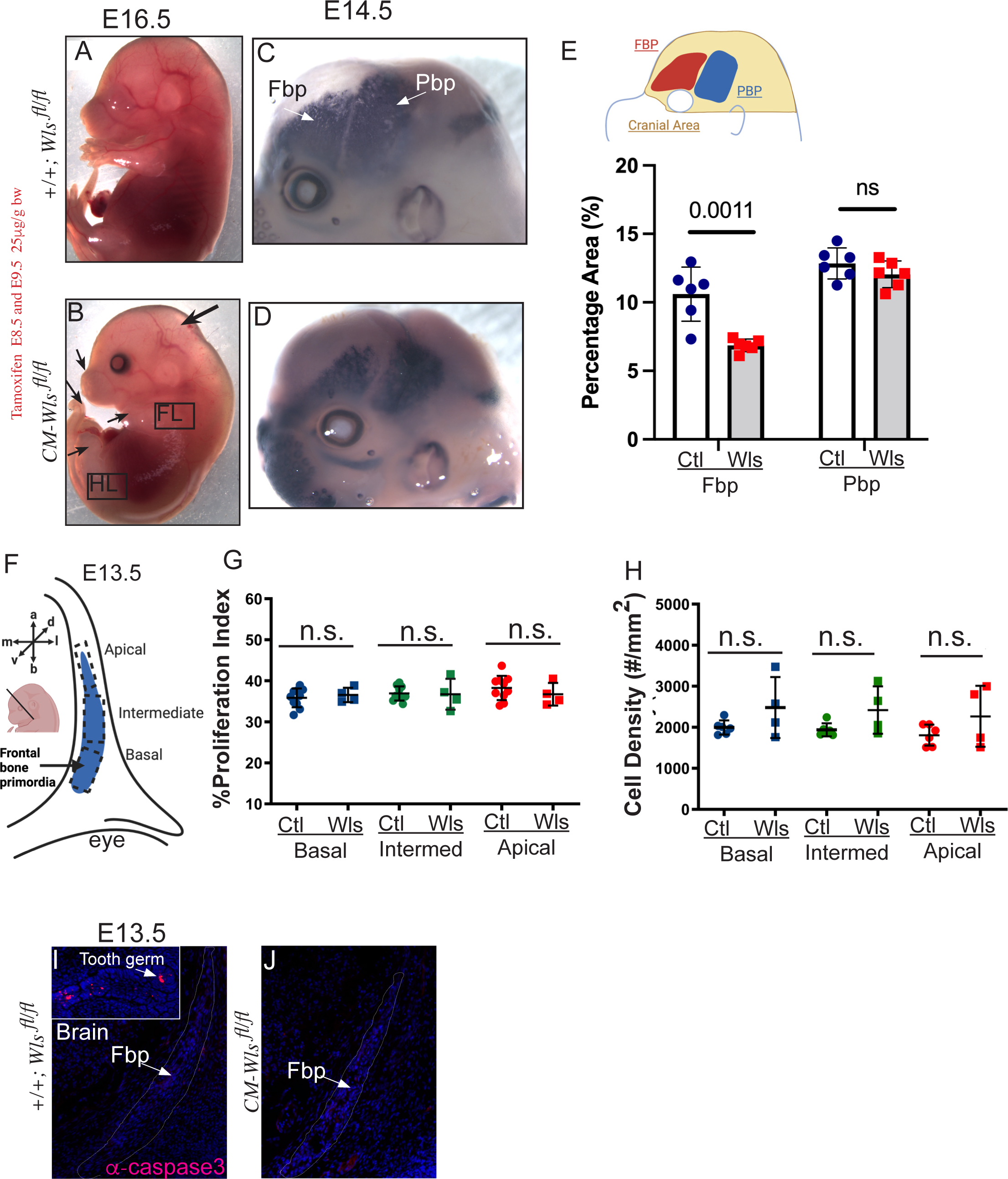
CM-*Wls* mutants have craniofacial extension defects without perturbing cell proliferation and survival. A, B) Wholemount view of control and CM-*Wls* mutants showing diminished extension of the fronto-nasal process, limb, and tail with a domed head. C-E) Frontal (Fbp) and parietal bone primordia (Pbp) stained with alkaline phosphatase in wholemount showing diminished frontal bone area normalized to cranial area. F) Schematic showing regions of interest in the frontal bone primordia (blue) at E13.5. G, H) Percent proliferation index by EdU incorporation for calvarial osteoblast proliferation and cell density in regions of interests were not significantly perturbed in CM-*Wls* mutants. E) Cell survival as assayed by activated caspase 3 expression by indirect immunofluorescence. Positive control in the tooth primordia. All control and mutant wholemount images are show at the same magnification.

**Supplemental Figure 4:**
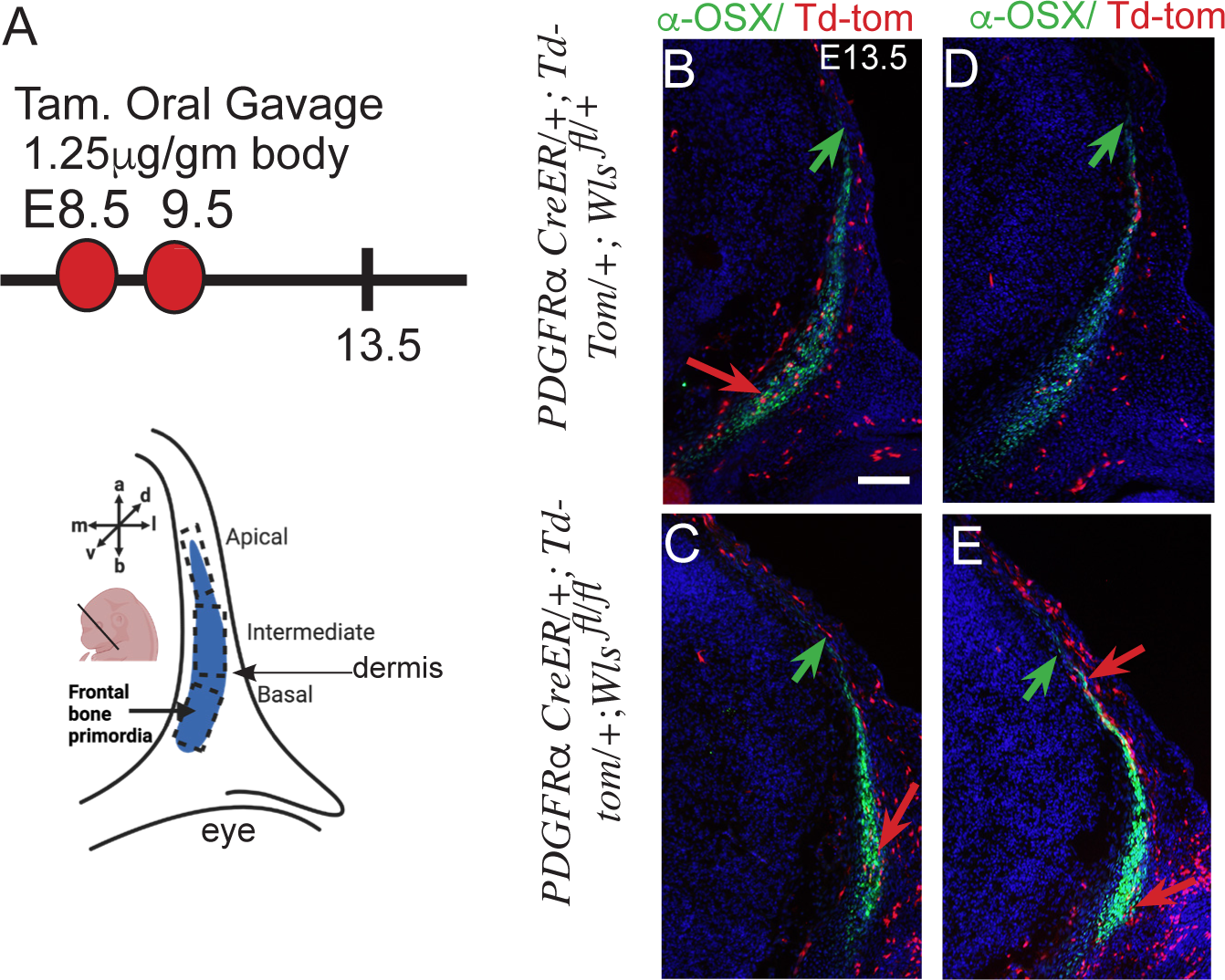
CM-*Wls* mutant cells can contribute to OSX+ frontal bone primordia. A) Schematic of the low dose tamoxifen gavage regimen to generate mosaic CM-*Wls* mutant cells in the SOA mesenchyme. B-E) Low dose tamoxifen leads to mosaic labeling of R26Td-Tomato in SOA mesenchyme in control and CM-*Wls* mutants at E13.5. Td-Tomato^+^ cells are visible in the OSX^+^ frontal bone primordia and the adjacent dermis in two different representative control and mutants (red arrows) (n=5 each). The apical position of the fbp is marked with green arrows. Scale bar =100 microns.

**Supplemental Figure 5:**
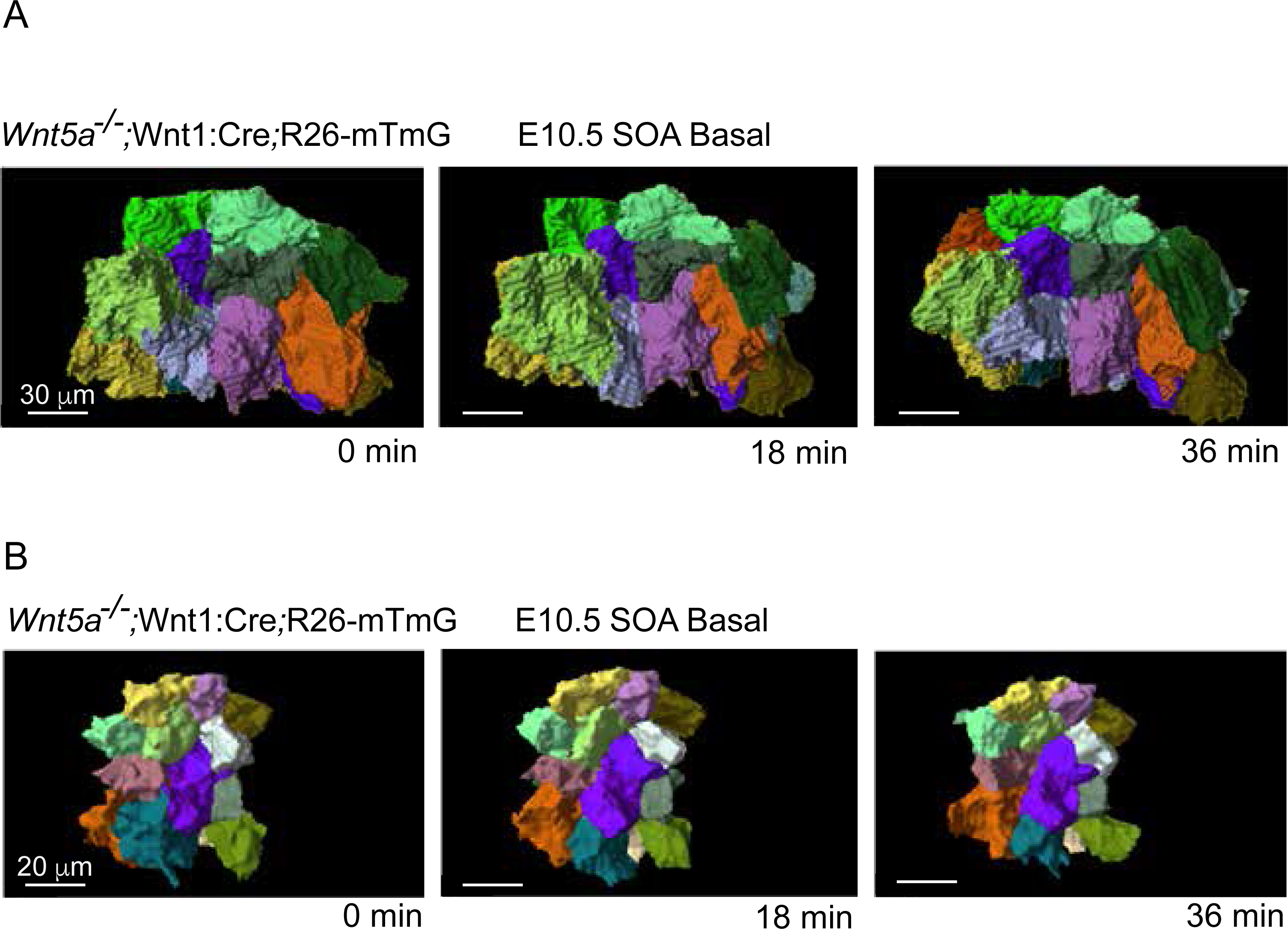
A,B) Two further examples of volume-rendered mesenchymal cells within the basal supraorbital regions of separate E10.5 *Wnt5a^-/-^;*Wnt1:Cre;R26:mTmG embryos. Despite cell shape fluctuations, mesenchymal cell neighbor positions remain stable.

**Supplemental Figure 6:**
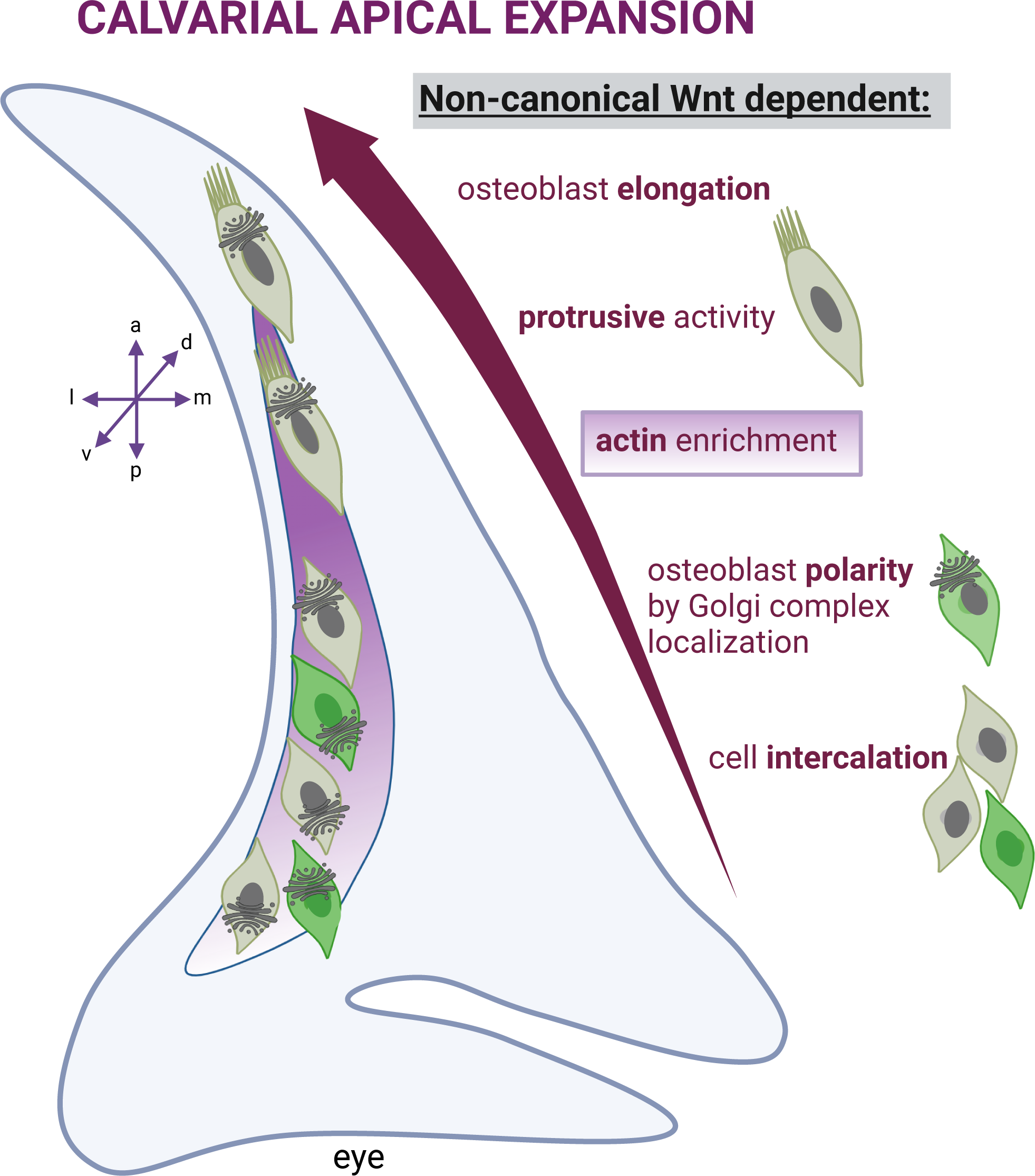
Summary schematic of the role of non-canonical mesenchyme Wnts in morphogenetic behaviors of calvarial osteoblasts.

## METHODS

### Mouse strains

Analysis was performed using the following mouse strains from Jackson Laboratory: mTmG Gt(ROSA)26Sortm4 (JAX Stock# 007676), (ACTB-tdTomato-EGFP)Luo/J)) (JAX Stock# 007576), 129S4.Cg-Tg(Wnt1-cre)2 Sor/J (JAX Stock #022137), CAG::myr-Venus (JAX stock# 16604528)*, Wnt5a^+/-^* (JAX stock# 10021340), *PdfgrαCreER* (JAX stock #018280) (Rivers et al., 2008), R26R-β-gal (JAX stock# 022616), R26TdTomato (JAX stock# 007905), and *Wntless^fl/fl^* mice (Carpenter et al., 2010) (gift of Richard Lang), were used in this project. All mouse lines were outbred to CD1 and genotyped as previously described for *PDFRαCreER/+*;*Wntless^fl/fl^*embryos, *PDFRαCreER/+; Wntless^fl/fl^*males and R26R/R26R; *Wntless^fl/fl^* females were time mated at night. After vaginal plugs were identified in the morning, days were counted by embryonic (E) 0.5. To induce CreER recombination, Tamoxifen at either 25μg or 0.125μg Tamoxifen/g body weight dose (Sigma T5648) was given by oral gavage twice to pregnant dams carrying E8.5 and E9.5 embryos. Tamoxifen was dissolved in corn oil and gavaged in the late afternoon of the designated day. For each experiment, a minimum of three controls and three mutant mice from at least 2 different litters were used. Case Western Reserve University Institutional Animal Care and Use Committee and Animal Care Committee of Hospital for Sick Children approved all animal procedures in accordance with AVMA guidelines (Protocol 2013–0156, Animal Welfare Assurance No. A3145-01).

### Live time-lapse light sheet microscopy and live time-lapse confocal microscopy

Three-dimensional (3D) time-lapse microscopy was performed on a Zeiss Lightsheet Z.1 microscope. Embryos were suspended in a solution of DMEM without phenol red containing 10% rat serum and 1% low-melt agarose (Invitrogen) in a glass capillary tube. Once the agarose had solidified, the capillary was submerged into an imaging chamber containing DMEM without phenol red, and the agarose plug was partially extruded from the capillary until the portion containing the embryo was completely outside of the capillary. The temperature of the imaging chamber was maintained at 37° C with 5% CO_2_. Images were acquired using a 20X/1.0 objective with single-side illumination, and a *z*-interval of 0.479 µm that was automatically calculated based on the numerical aperture (1.0) of the objective. Images were acquired for 2-3 hours with 3 minute intervals. Fluorescent beads (Fluorospheres 1µm, Thermo Fisher, 1:10^6^) were used as fiducial markers for 3D reconstruction and to aid in drift-correction for cell tracking. Multi-view processing was performed with Zen 2014 SP1 software to merge the 3 separate views and generate a single 3-dimensional image. Further analysis and cell tracking were performed using Vision4D (Arivis) software. Spinning disk confocal microscopy in whole embryos was performed to visualize SOA cranial mesenchyme cells using the Quorum WaveFX-X1 (Quorum technologies) as described previously (Tao et al., 2019).

### Membrane segmentation and 3D cell neighbour counting

Timelapse 3D datasets of cell membranes were processed with the ImageJ macro Tissue Cell Segment Movie (kindly provided by Dr. Sébastien Tosi from the Advanced Digital Microscopy Core Facility of the IRB Barcelona) to generate membrane segments of embryos expressing the cell membrane reporter, *ROSA26;mTmG* prior to analysis with Imaris 9.0 software (Bitplane). Surface objects were created using Imaris and, using the “Add New Cells” function, cell boundaries from a region of interest were detected using the local contrast filter type and fluorescence intensity/quality thresholds. Cell tracking was performed using the autoregressive motion algorithm. Analysis was performed on 3 separate areas in basal and apical regions of the supraorbital arch over 2 independent experiments for each condition.

### Strain analysis

Tracking of tissue deformation was performed over 16 time steps in the rostrocaudal-mediolateral plane of time-lapse light sheet movies of the supraorbital arch. For each sample, tracking was performed for ∼50 points by image correlation of a 24 x 24 pixel box centered on each point in subsequent images. Image correlation was done using a subpixel registration algorithm explained in (Guizar-Sicairos et al., 2008). Once points were tracked, we applied Delauney tesselation to generate a triangular mesh from the initial positions representing the initial tissue shape. We then calculated strain in the horizontal and vertical directions for each triangular element and assigned to each point the area weighted average of elements including that point. Custom tool used for deformation tracking is available at [https://github.com/HopyanLab/Strain_Tool].

### Histology, Immunohistochemistry, and staining

Heads of E12.5–14.5 embryos were drop-fixed in 4% paraformaldehyde (PFA) for 30–45 min, respectively, at 4°C and cryopreserved as previously described(Atit et al., 2006). Embryos were cryosectioned at 14 μm in the coronal plane of the frontal bone primordia. For immunofluorescence cryosections were dried at room temperature for 10 minutes, washed in 1X PBS and blocked in 10% specie-specific serum with 0.01% Triton for 1 hour . Primary antibodies were diluted in blocking buffer and slides were incubated overnight at 4 °C. The next day, the cryosections were washed in 1X PBS, incubated with species-specific fluorescent secondary antibodies (below) for one hour at room temperature, washed with DAPI 0.5 μg/mL, and mounted with Fluoroshield (Sigma F6057). The following primary antibodies were used for immunofluorescence experiments: rabbit anti-Osx (1:2000 or 1:4000; Abcam ab209484), rabbit anti-Caspase 3 (1: 250; Abcam ab13847), Goat anti-Runx 2 (1:100; R&D Systems AF2006-SP) and mouse anti-GM130 (1:100; Thermo Fisher Scientific BDB610822). Appropriate species-specific Alexa-fluor secondary antibodies were used (Alexa 488, 1:500, Thermo Fisher A32790 and Alexa 594, 1:800 Thermo-Fisher A11012). For immunohistochemistry staining against GM130 (BD Dickinson), Vector M.O.M kit (BMK2202) was used for blocking before incubating with primary antibody, Diva-Decloaker (Biocare) was used for heat mediated antigen retrieval for 15 min, and Vector ABC elite kit was used for tertiary amplification followed by staining with DAB. For Concanavalin A membrane staining, sections were dried at room temperature, rinsed in Hanks buffered salt solution (HBSS), stained in 10ug/ml Concanavalin-A (Thermo Fisher C11253) diluted in Hanks balanced buffered solution for 30 minutes at room-temperature. Sections were rinsed in HBSS and counterstained with DAPI (1:2000) and mounted with Fluoroshield. Phalloidin-488 conjugate (1:1000) was used to visualize filamentous actin staining by incubating samples for 30 min at room temperature before rinsing cells in 1X PBS and counterstaining with DAPI (1:2000) and mounting with Fluoroshield. Alkaline phosphatase(AP) staining was performed as previously described(Ferguson et al., 2018). AP stained sections were imaged on the Hamamatsu Nanozoomer S60 Digital Whole Slide Scanner.

### Cell proliferation, cell density, cell survival

Mice were administered 250 μg EdU in PBS/10 g mouse weight by intraperitoneal injection one hour prior to sacrifice. Embryos were then collected and prepared for cryopreservation as previously stated, EdU staining was conducted using the Click-iT Plus EdU kit (Thermo Fisher Scientific, C10639) using the standard protocol suggested by the manufacturer. The percent EdU^+^ positive calvarial osteoblasts/total DAPI^+^ nuclei was calculated in the coronal sections of frontal bone primordia that was identified by morphology of condensed nuclei. Three to four different sections per embryo were used for analysis using ImageJ/Fiji, Adobe Photoshop and Cell Profiler. First, regions of interests (ROI) in the apical, intermediate and basal regions of the frontal bone primordia were generated in Fiji. Then, images were cropped in Adobe Photoshop and analyzed in Cell Profiler using the following pipeline: a. split each image into two channels: blue and red, b. smooth using a Gaussian filter, c. uneven background illumination was calculated using a block size of 22 for both channels, d. global two class threshold strategy was employed using the Otsu method for objects with a diameter of 10-30 (EdU positive cells), and 10-15 (DAPI positive cells). The number of EdU positive and DAPI positive cells were exported as a Microsoft Excel spreadsheet.

### Histomorphometrics and imaging

For length-width ratio of Alkaline phosphatase stained frontal bone primordia in coronal plane sections was obtained by doing a point to point measurement for the length and widest width in Image J/Fiji (Schneider et al., 2012). Length-width ratio was calculated from one histological section of the frontal bone primordia from E13.0 and E14.5 (n=3-4 wild type embryos). Similarly, length-width ratio of individual calvarial osteoblasts cells and nuclei was obtained regionally basal, intermediate or apical region of interest in the frontal bone primordia after immunostaining for RUNX2 or OSX. Length-width ratio of calvarial osteoblasts was obtained from OSX+ cells co-stained with Concavalin A lectin for membranes at E13.5 (n=4 wild type embryos). The length-width ratio of Runx2^+^nuclei was obtained at E12.5 (n=4-5 embryos).

For quantifying the normalized apical extension of the frontal bone, frozen sections of embryonic mouse heads in the coronal plane were stained for alkaline phosphatase as described above. Images from the Hamamatsu Nanozoomer were opened on the NDPI viewer software. Using a fixed zoom, the baso-apical length of the frontal bone primordia was measured using the freehand line tool and converted to microns. Similarly, the cranial length was measured from the eyelid to the midline of the brain. The percent apical extension was calculated as the ratio of length of the frontal bone to the cranial length. Frontal and parietal bone primordia area was obtained from wholemount alkaline phosphatase stained embryos. Individual bone primordia area measurements was normalized to cranial area in Fiji.

GM130+ Golgi complex to nuclei angle was calculated in the RUNX2^+^ or OSX^+^ expression domain of the frontal bone primordia in the coronal plane. The image was oriented so that the eyes were along the x-axis (0 degree) and the midline of the brain/apex along the y-axis (90 degrees). The Golgi complex-nuclei angle was measured relative to the center of the nuclei manually using Image J/ Fiji software. Approximately, 25-30 cells in apical, 35-45 intermediate and 50 cells in the basal region of the frontal bone primordia were randomly selected from controls and CM-*Wls* mutants were selected in MS Excel (n=4 embryos, 3-4 sections/embryo).

All widefield images were captured using the Olympus BX60 microscope, Olympus DP70 digital camera using DC controller software. Images were processed in Adobe Photoshop 2022, Image J /Fiji (Schneider et al., 2012) and assembled on Adobe Indesign or Illustrator 2022.

### Statistical analysis

All graphs were created using Prism 9 (GraphPad Software: https://www.graphpad.com/scientific-software/prism/). All statistical analysis was conducted on Prism 8, using an unpaired, two tailed, *t-*test with Welsh’s correction. Circular distribution of Golgi-complex nuclei angles was analyzed by Kolmogrov-Smirnov test (Apte and Marshall, 2013) . Data that is not significant is denoted by “n.s.”. The p-values for statistical tests in all figures are stated or represented as: * = p < 0.05, ** =p < 0.01, and *** = p <0.001.

## Acknowledgements

Thanks all the present and past members of the Atit Lab and Hopyan Lab for contributing ideas or effort to this project. Special thanks to Nyoka Lovelace and Mia Carr for help with animal husbandry and immunostaining, James Ferguson and L. Henry Goodnough for expression analysis of Wnt/PCP components, Yifan Zhai for frontal bone area analysis. Thanks to CWRU-bio shared instrumentation facility for access to microscopes. Schematics were made on biorender.com. This work was supported by NIH-NIDCR R01-DE18470 (RA), NIH-NIDCR R21-DE029348 (RA, SH), NIH-NIDCR F31 NRSA fellowship (TI), NIH-NIAMS T32 fellowship (JF), Case SOURCE fellowship (NP), Canadian Institutes of Health Research MOP 126115 (SH), Canadian Institutes of Health Research (168992) (SH), and the Canada First Research Excellence Fund/Medicine by Design (MbDGQ-2021-04) (SH).

